# Inferring recent demography from isolation by distance of long shared sequence blocks

**DOI:** 10.1101/076810

**Authors:** Harald Ringbauer, Graham Coop, Nick Barton

## Abstract

Recently it has become feasible to detect long blocks of almost identical sequence shared between pairs of genomes. These so called IBD-blocks are direct traces of recent coalescence events, and as such contain ample signal for inferring recent demography. Here, we examine sharing of such blocks in two-dimensional populations with local migration. Using a diffusion approximation to trace genetic ancestry back in time, we derive analytical formulas for patterns of isolation by distance of long IBD-blocks, which can also incorporate recent population density changes. As a main result, we introduce an inference scheme that uses a composite likelihood approach to fit observed block sharing to these formulas. We assess our inference method on simulated block sharing data under several standard population genetics models. We first validate the diffusion approximation by showing that the theoretical results closely match simulated block sharing patterns. We then show that our inference scheme rather accurately and robustly recovers estimates of the dispersal rate and effective density, as well as bounds on recent dynamics of population density. To demonstrate an application, we use our estimation scheme to explore the fit of a diffusion model to Eastern European samples in the POPRES data set. We show that ancestry diffusing with a rate of 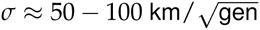 during the last centuries, combined with accelerating population growth, can explain the observed exponential decay of block sharing with pairwise sample distance.

---

There has been longstanding interest in estimating demography, as migration and population density are key parameters for evolution and ecology. Having a demographic model is essential for disentangling the effects of neutral evolution from selection, and hence crucial to understanding adaptation. Moreover, inference of demographic parameters is important for conservation and breeding management. As direct observations are often work-intensive and necessarily limited to short time scales, the increasing availability of genetic markers has spurred efforts to develop inference methods based on genetic data.

This work focuses on estimating demography by using pairwise shared long blocks of genome that originate from recent genetic co-ancestry. Specifically, we study so called IBD - blocks (identity by descent) in sexually reproducing populations, which are commonly defined as co-inherited segments delimited by recombination events on the path to the common ancestor (see Fig. 1). It has now become feasible to detect long regions of exceptional pairwise similarity from dense SNP or whole genome sequences (e.g. Gusev *et al*. (2009); Browning and Browning (2011)) and for regions longer than a few cM, the bulk of such regions mostly consists of a single IBD-block unbroken by recombination, at least when inbreeding is rare (Chiang *et al*. 2016). This yields novel opportunities for inferring recent demography, as one can study the direct traces of recent co-ancestry.

**Figure 1.**
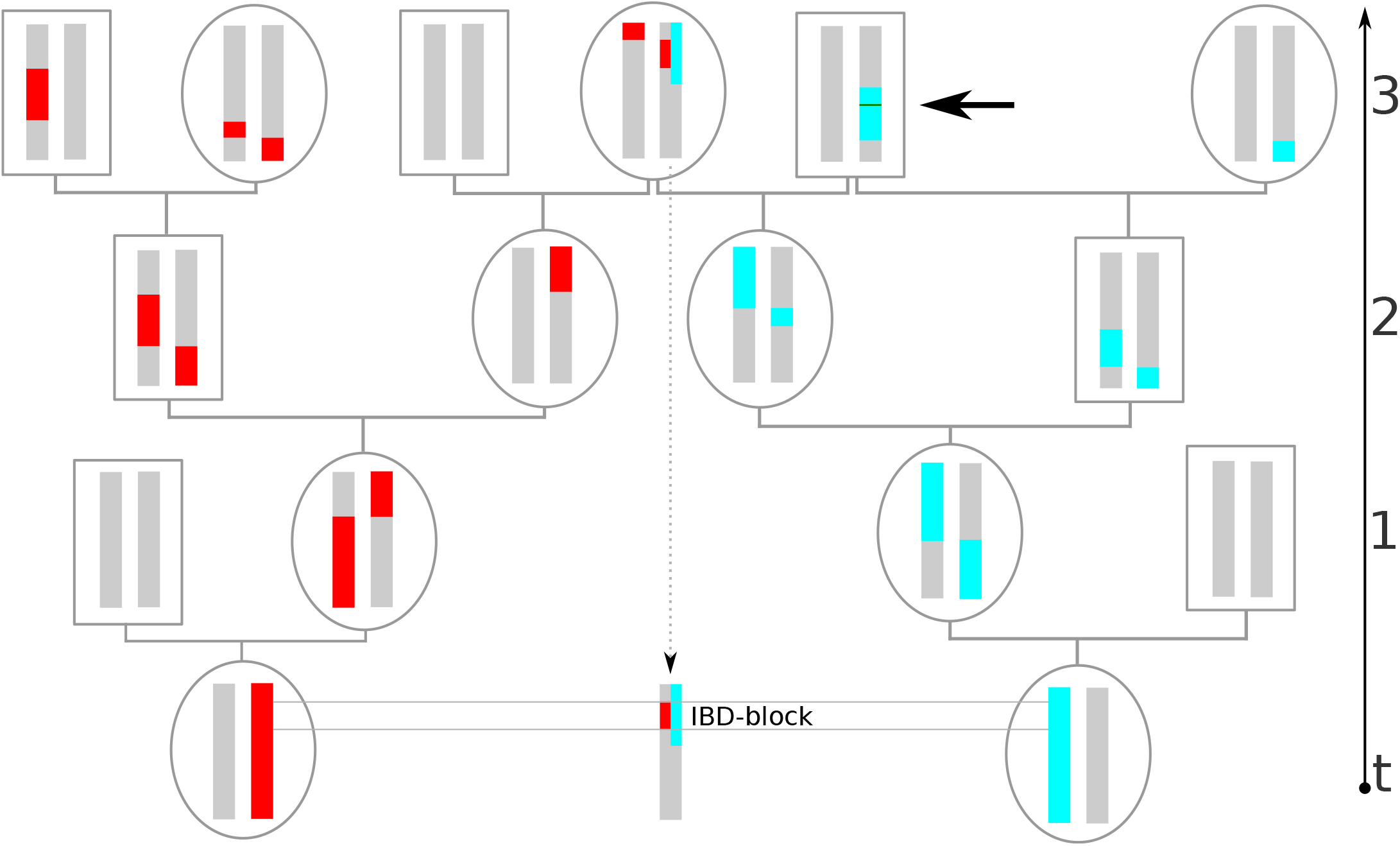
Example of an IBD-block co-inherited from a common ancestor three generations back. Going back in time, recombination splits up genetic ancestry (colored here red and blue) into blocks distributed among ancestors. If, as depicted here, such ancestral blocks overlap in a recent common ancestor, the intersecting stretch of genome will be shared and both individuals carry only few distinguishing mutations there. Here we define IBD-blocks to be delimited by any recombination events on the genealogical path to the most recent common ancestor. Thus, also recombination events that are fused again quickly by inbreeding loops, as depicted for the blue chromosome (thick arrow), delimit IBD-blocks. This recombination is not detectable in practice however, and the two adjacent IBD-blocks would be identified as one long IBD-segment.

Recombination can then be used as a clock, since the length of shared blocks contains information about their age: The longer the time to the most recent common ancestor, the shorter the expected IBD length, as recombination has had more chance to break up ancestral genetic material – the probability that a block of a given map length is not hit by recombination decays approximately exponentially with time back. Long IBD-blocks hence originate mostly from very recent co-ancestry. Analyzing them yields insight into very recent times and is complementary to the analysis of short identical segments, which are informative about deeper times scales (Li and Durbin 2011; Harris and Nielsen 2013).

Recent work has begun to explore this potential for inference of recent demography: Sharing of long blocks between pairs of populations can be used to infer the distribution of recent coalescence times (Ralph and Coop 2013), and fitting deme and island models can yield information on recent population sizes (Palamara *et al*. 2012; Browning and Browning 2015) and migration patterns (Palamara and Pe’er 2013).

Here, we focus on a pattern of isolation by distance of IBD-blocks within populations extended in two dimensions with local migration. For such populations, the classical Wright-Malecot formula describes an increase of mean pairwise genetic diversity with increasing geographic separation, a classical isolation by distance pattern (Wright 1943; Malécot 1948). Several demographic inference methods utilize this as signal, typically by fitting the predicted pattern of increasing pairwise F_ST_ with geographic distance, to infer parameters of recent demography (e.g. Rousset (1997, 2000); Vekemans and Hardy (2004)). However, such schemes based on pairwise diversity have fundamental limitations. Importantly, these methods can usually only infer the so called neighborhood size, the product 4*πDσ*^2^ of the dispersal rate *σ*^2^ with effective density *D*; separate estimation of these important parameters is usually not feasible, as the underlying signal is based on a relatively short-term equilibrium between local drift and dispersal, and typically mutation rates are too low to provide significant additional information on these short time scales (Barton *et al*. 2013). Moreover, theoretically expected isolation by distance patterns on small scales, that built up over relatively recent times (Leblois *et al*. 2004) can be severely confounded by deeper, often unknown ancestral patterns (Meirmans 2012).

This work expands on Barton *et al*. (2013) to overcome these problems: As in the derivation of the Wright-Malecot formula, they use spatial diffusion to model the spread of ancestral genetic material back in time to derive first analytical formulas for block sharing depending on pairwise sample distance and block length. Based on these theoretical results, they observed that analysis of shared IBD-blocks would in principle allow one to use recombination as a clock to estimate dispersal and past density separately, very robust to confounding by ancestral structure.

Here, we introduce a practical inference scheme based on this idea, that fits the underlying demographic model by maximizing a composite likelihood, similar to Ralph and Coop (2013). For this, we first expand the analytical formulas of Barton *et al*. (2013) to explicitly describe the expected number of shared blocks of a certain block length for a given sample pair. Similarly, we use geographic diffusion of ancestry in our model, and we additionally describe the effect of recent population density changes. Since formally using a geographic diffusion approximation is not always well justified (Felsenstein 1975; Barton *et al*. 2002), we here also demonstrate that the derived formulae closely match simulated IBD-block sharing using several standard models of geographically extended populations.

The resulting inference scheme can estimate the dispersal rate and past population density separately. It can be also used to infer patterns of global population density growth or decline, and in contrast to existing methods for inferring recent population size changes based on IBD-blocks (Browning and Browning 2015), it takes the geographic spread of ancestry into account. We test the estimation scheme on simulated IBD-sharing data, which empirically demonstrates its power to recover demographic parameters from patterns of block sharing.

Recently, Baharian *et al*. (2016) have independently derived similar formulas for block sharing under the diffusion approximation for a constant population density, and applied their result to IBD-sharing between human North American samples. They estimated dispersal and population density by fitting a formula for the expected sum of lengths of all IBD-blocks longer than a threshold, where they binned sample pairs according to pairwise geographic distance. In contrast to this regression, our composite likelihood scheme additionally incorporates information from the exact lengths of shared blocks, and directly fits pairwise block sharing without first averaging over distance bins. As we show below, the likelihood framework also allows one to readily include error estimates for block detection, such as limited detection power or wrongly inferred block lengths, as often unavoidable when calling IBD-segments from genotype data (Browning and Browning 2012; Ralph and Coop 2013). Importantly, our inference scheme also incorporates recent population decline or growth, which is likely important for the analysis of human data.

The formulas and the inference scheme describe IBD-blocks as defined above, where every recombination event on the path to the common ancestor is assumed to delimit an IBD-segment. However, adjacent IBD-blocks, such as produced by chance or quick inbreeding loops (Fig. 1), are in practice detected as one single long IBD-block. Here we demonstrate with simulations that this effect does not markedly influence inference as long as local inbreeding is not unreasonably common, and only starts to significantly confound estimates when the density of individuals measured in dispersal units is about one.

At the moment, the largest IBD-block data sets including accurate information on sample locations are available for humans. To showcase a practical application of the inference scheme, we apply it to a subset of the POPRES dataset, which Ralph and Coop (2013) have previously analyzed for sharing of long IBD-blocks. We use their publicly available data and detailed estimates of block detection errors, which we included into our inference scheme. Human demography is likely very complex, but the diffusion model provides a good fit to the data that allows us to draw conclusions about the extent of human ancestry spread during the last centuries in continental Europe. We also detect a clear signal of a rapidly increasing population density, which stresses the importance for accounting for rapid population growth when analyzing human IBD-sharing.

## Materials and Methods

### The model

To describe block sharing in two spatial dimensions with local migration we use two basic model assumptions, which should approximate a wide range of scenarios. Obviously the true demographic history of a species will often be more complex than any such simple model. Thus the aim is not to have a mathematically rigorous model, which is often formally problematic (Felsenstein 1975; Nagylaki 1978) and only holds exactly in very specific settings, but an accurate approximation that captures general patterns that can be used for robust inference of basic demographic parameters. In the following we outline these two central modeling assumptions.

#### Poisson recombination

We approximate recombination as a homogeneous Poisson process, i.e. crossover events are assumed to occur at a uniform rate along a chromosome. This implies that genetic distances are measured in map and not physical units. Throughout this work the unit of genetic distance will therefore be the Morgan, which is defined as the distance over which the expected average number of intervening chromosomal crossovers in a single generation is one. Small scale processes such as gene conversion are not captured by the Poisson approximation, but for the large genomic scales of typically several cM (1 Morgan = 100 cM) considered here this can be neglected (Lynch *et al*. 2014). We also ignore the effect of interference, which is reasonable when describing the effects of recombination over several generations. Moreover, the female and male recombination rate can be markedly different. For the purpose here we can use the sex averaged rate 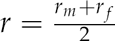: In every generation loci on autosomes have an equal chance to trace back to a female or male ancestor, thus the female and male Poisson process together are described by a single Poisson process with the averaged recombination rate. Generalizing this line of thought, any individual differences in map length are modeled here with a single Poisson process with the population averaged rate.

#### Diffusion approximation

Following a long tradition of modeling individual movement in space with diffusion (Fisher 1937; Wright 1943; Malécot 1948; Nagylaki 1978), we approximate the spatial movement of genetic material back in time by a diffusion process: The position of ancestral material at some time *t* in the past is the sum of migration events until then, which are often correlated only on small time scales. Therefore, by the central limit theorem, the probability density for the displacement of a lineage can be approximated by a Gaussian with axial variance *σ*^2^*t* that linearly increases back in time *t* (Fig. 2). Importantly, this is independent of details of the single-generation dispersal kernel, provided its variance is finite. The diffusion approximation typically needs several generations of migration events to become accurate, whereas it often also does not hold over very long timescales when large scale events such as colonization become important. However, on the recent to intermediate timescales relevant for the sharing of long IBD-blocks, it seems plausible that diffusion often accurately describes the spread of ancestry – as long as populations have reached a local equilibrium (Barton *et al*. 2002, 2013). If consecutive single generation dispersal events are uncorrelated, *σ*^2^ is simply the average squared axial parent-offspring distance. Even in face of small scale spatial or temporal correlations between dispersal events one can still hope to model the spread of ancestry by diffusion (Robledo-Arnuncio and Rousset 2010). Then *σ*^2^ has to be interpreted as a parameter describing the rate of spread of ancestry back in time (Barton *et al*. 2002), which can be markedly different from the single generation squared axial parent-offspring distance.

**Figure 2.**
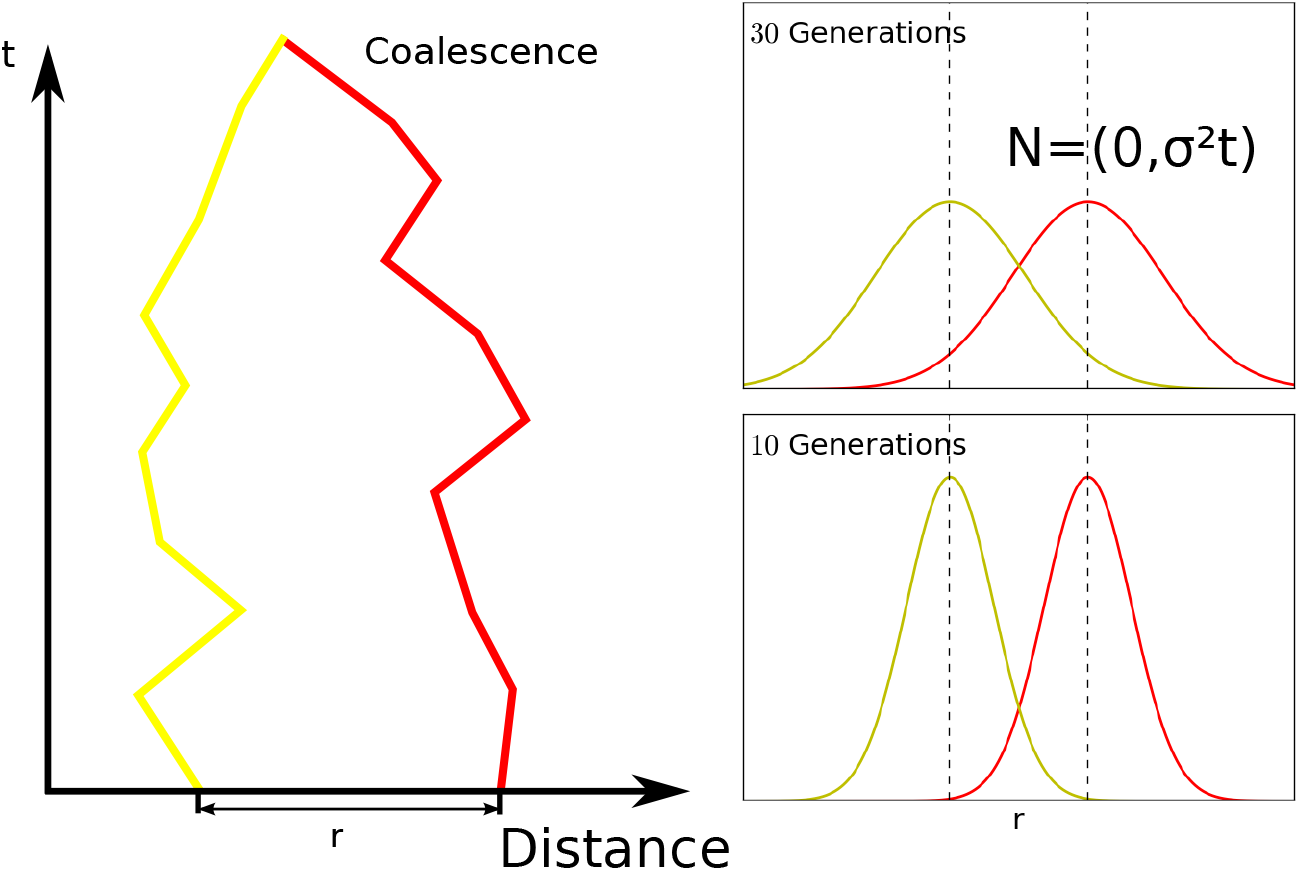
Diffusion model in one spatial dimension. Left: One realization of movement of ancestry of two homologous loci initially separated by distance *r* back in time. Right: In our model, the probability density function of having moved distance ∆*x* at time *t* generations back spreads out as a Gaussian *N*(0, *σ*^2^*t*) with linearly increasing variance *σ*^2^*t*.

This diffusion approximation implicitly assumes neutrality for most parts of the genome, as selection can also influence the position of ancestral lineages. However, on the relatively short timescales considered here this is expected to significantly affect the geographic position of ancestral lineages only in the case of ongoing strong sweeps.

Here we need to describe the distance between pairs of lineages of homologous loci. We approximate this by adding the movement of two single lineages, which yields a Gaussian with twice the variance 2*σ*^2^*t* for the separating geographic distance at time *t* back. This approach of folding two independent random walks implicitly assumes that movement of a pair of lineages is independent. Typically correlations of movements are of geographically very local nature, e.g. in case of density fluctuations or local barriers, and often average out when viewed on larger scales (Barton *et al*. 2002). Thus this folding should be a sound approximation when describing the chance that ancestral genetic material of initially spatially well separated genetic material comes close. It furthermore ignores that once coalesced, lineages remain at pairwise distance 0. However, this can be safely neglected as long as pairwise coalescence is rare – as usually the case when only a small fraction of genetic material coalesces in the very recent past.

Felsenstein (1975) observed that any spatial model that incorporates Poisson offspring distribution and independent migration following a fixed dispersal kernel (as implicitly used by Malécot (1948)) will inevitably result in clumping of individuals, thus making any such model biologically unrealistic. Here, we use diffusion to trace genetic material backwards in time and not forwards. While modeling movements of pairs of lineages to be uncorrelated, we do not invoke this paradox in its original form related to clumping: Following a pairwise coalescent process back with diffusion does not implicitly assume that individuals have independent number of offspring.

### IBD-sharing in the model

Using similar assumptions, Barton *et al*. (2013) calculated the probability that two individuals a certain distance apart share an IBD block longer than a minimum length starting from a specified locus. For this, the Wright-Malecot formula can be applied directly by replacing mutation with recombination. This was intended to be a proof of principle; for practical inference from IBD-blocks, formulas describing the total number of shared block of a specific given length L are advantageous. For this one has to take a different route, which we begin with a calculation following Ralph and Coop (2013):

#### IBD-Blocks of age *t*

For a given pair of samples, we partition *N*_*L*_, the number of shared blocks of map length L into 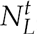, the number of blocks coalescing at time *t* ago. Here and throughout this paper, such terms are always understood as a density with respect to block length and time, which integrated over an area [*L*_1_, *L*_2_] resp. [*t*_1_, *t*_2_] yields the expected number within it. Using the linearity of expectations gives:

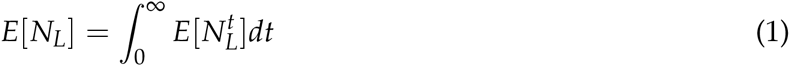

Following the ancestry of two chromosomes back until time *t* before the present, a change of genealogy only occurs when there is a recombination event somewhere along the lineage. Between these discrete jumps, genetic material can be traced as a single locus. Due to the independence of recombination from the underlying genealogy, we can use this to further split 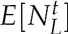 into the product of the expected number of blocks of length *L* obtained by splitting the two chromosomes according to Poisson recombination over time *t* with the probability that a single locus coalesces time *t* ago. We denote the first factor by *K*_*t*_(*L*), the second, commonly known as the coalescence time distribution, by *µ*(*t*):

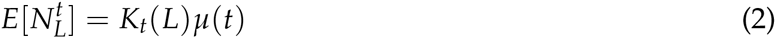

#### Number of candidate blocks

Under our model assumptions, the position of all recombination events on genetic material of two independent chromosomes traced back until time *t* is given by a Poisson process with rate 2*t*. The expected number of all block pairs overlapping at an intersection length *L* can then be calculated straightforwardly: A small region of map length Δ*L* is hit with probability 2*t*Δ*L* by a recombination event, and the probability that a region of length *L* does not recombine follows a exponential distribution exp(-2*Lt*). For chromosomes of map length *G*, using the linearity of expectations and summing over all possible start sites yields the expected total number of blocks of length *L*: 
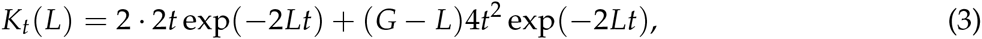
where the first term describes blocks starting at either edge and the second term fully interior blocks, which require two delimiting recombinations. Neglecting chromosal edge effects (*G* >> *L*) this is approximated by 
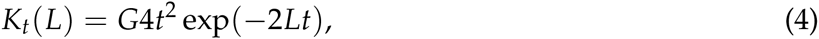
which we will use for deriving an approximate formula capturing the qualitative behavior of mean IBD-sharing. The slightly more complex full result including edge effects used for inference is derived analogously (see Appendix B).

#### Single locus coalescence probabilities

The probability *µ*(*t*) that two homologous loci have their last common ancestor time *t* ago will depend on their pairwise sample distance *r,* and the parameters of the demographic model. Here we proceed as in Barton *et al.* (2002) and approximate the probability of recent coalescence as the product of the probability of the pairwise sample distance being 0 and a rate of local coalescence which, following previous literature (e.g. Barton *et al.* (2013)), we shall denote here by 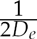. In Appendix *A* we justify this approximation and also give a formal definition of this so called effective density *D*_*e*_. In order to describe a globally growing or declining population, which is particularly important for the human case studied here, we let *D*_*e*_ depend on time *t*. This yields:

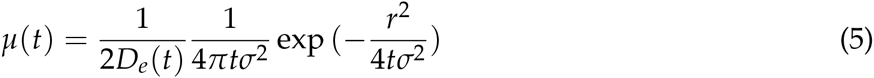

#### Full formula

Substituting into formula Eq. 4 and Eq. 5 into Eq. 2 gives:

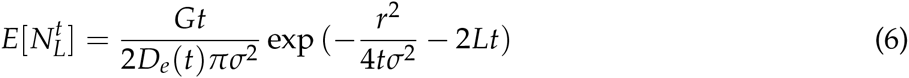

For the total number of expected shared blocks we have to integrate over all possible coalescence times *t*. For the class of power density functions, 
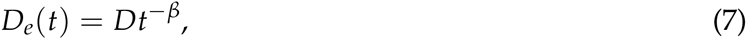
the integral yields explicit formulas. Here *D* > 0, and 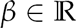 can be interpreted as a growth parameter. The important case of *β* = 0 models a constant population density, while *β* > 0 and *β* < 0 describe populations with growing or declining density respectively. This class of functions has been used to fit human demographic growth (Von Foerster *et al.* 1960), but importantly, more complex density functions can be fitted as linear combinations of such terms (including polynomials for the special case 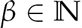). Due to the linearity of the integral this yields explicit formulas too, which numerically greatly simplifies inference.

When using a single power term only, with *β* > 0 the density approaches infinity for *t* = 0, which corresponds to a negligible chance of coalescence at the present. This is not as limiting as it looks on first glance: Since genetic material of a pair of samples several dispersal distances away need some time to reach each other, one effectively fits density on intermediate time scales (see also Fig. 11).

Performing now the integral of Eq. 1 yields the main result:

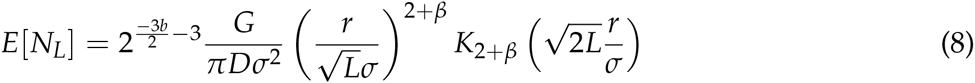

Integrating this with respect to block length gives the expected number of shared blocks longer than a threshold length *L*_0_: 
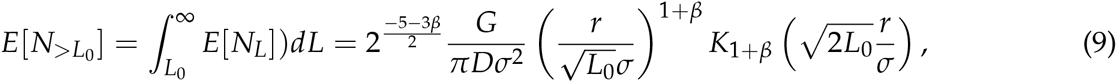
where *K*_*γ*_ denotes the modified Bessel-function of the second kind of degree *γ* (Abramowitz and Stegun 1964). These results are analyzed qualitatively in the discussion section, and Fig. 3 depicts their accuracy on simulated data. For the special case of a constant population size (*β* = 0), one arrives at an equivalent formula independently derived by Baharian *et al.* (2016), who describe the expected fraction of genome shared through segments within a given length range instead of the expected number of shared blocks. The approach outlined here also allows us to incorporate chromosomal edge effects (see Appendix B).

**Figure 3.**
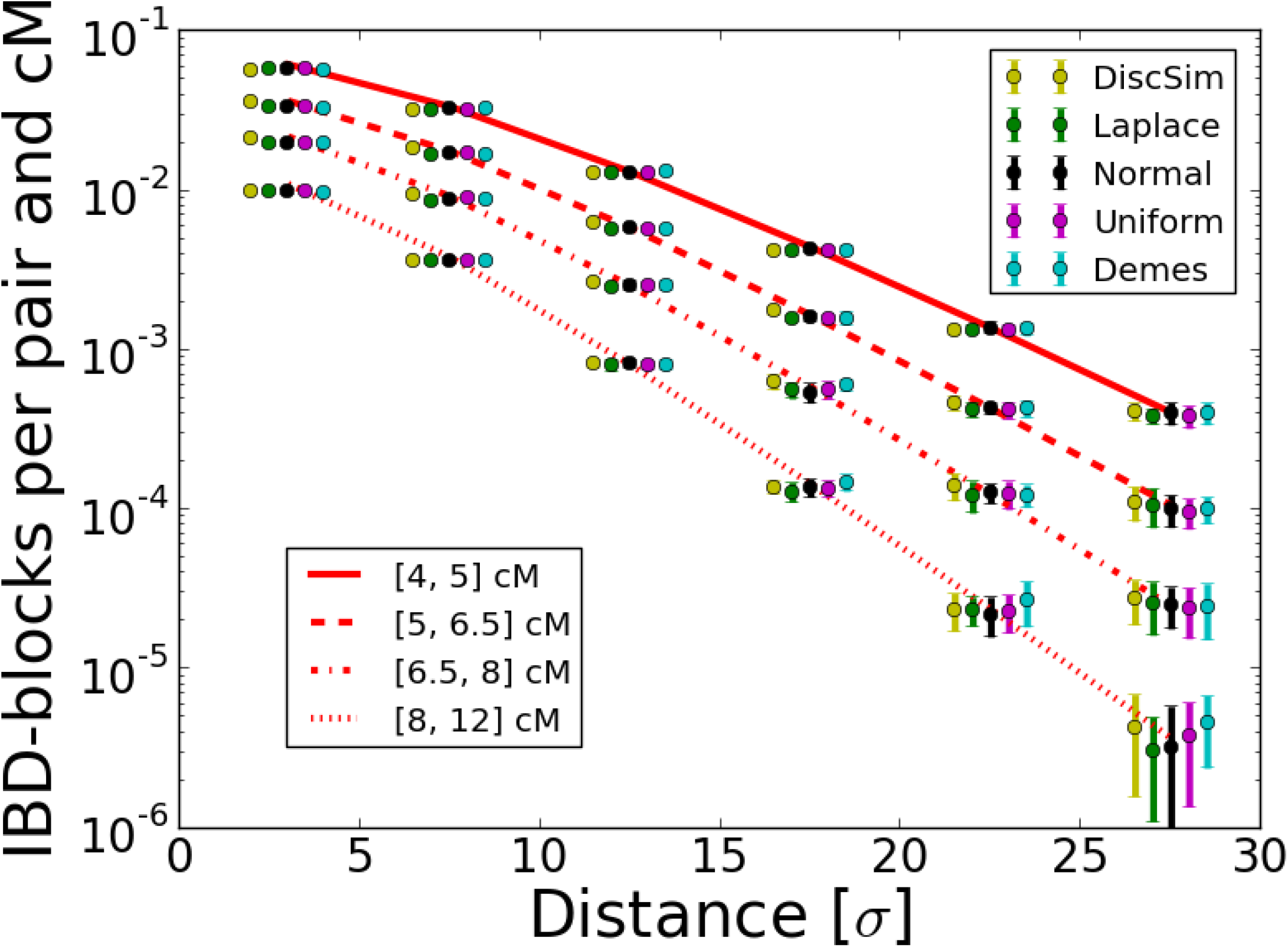
Simulated IBD-block sharing compared to theory. We show values normalized to give rates per pair and cM. Theoretical expectations are calculated for each length bin via Eq. 9. For five models, we kept population density constant at *D*_*e*_ = 1, with a dispersal rate *σ* = 2 on a torus of size 180 and simulated IBD-sharing between 150 cM chromosomes spread out on a sub-grid in nodes 2 distance units apart (For full set of specific simulation parameters see Supplementary Text 2). For every model we ran 20 replicate simulations. In the figure, distances are measured in dispersal units (so that *σ* = 1) and error bars depict the estimated standard deviations for each bin among the 20 runs to visualize uncertainty of estimates. Dots are spread out for better visualization around their original position (middle dot for normal dispersal)

### Inference scheme

In order to learn about recent demography one has to fit observed block-sharing between a set of samples to the here derived formulas (Eq. 8). Here we use a likelihood method: We approximate the likelihood function 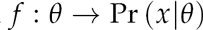 of the observed data *x* for a given set of parameters *θ* (here *σ, D*, *β*) with a composite likelihood 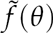, and use standard numerical optimization techniques to find the maximizing estimates 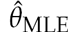.

#### Assumption of independence

Working with expected values is useful in the derivation because of their linearity, but a likelihood-scheme requires explicit probabilities for sharing IBD-blocks. Here we take the same approach as Ralph and Coop (2013): Different shared blocks among pairs are assumed to be the outcome of independent processes.

This is obviously an over-simplification: Block-sharing can be correlated along chromosomes and also among different sample pairs, due to initially shared movement of genetic material. Taking all these correlations into account would go beyond the simple pairwise diffusion model. However, for the purpose of fitting the assumption of independence is not too limiting. Maximizing the likelihood of actually correlated observations (composite likelihood) is a widely used practice in inference from genetic data (e.g. Fearnhead and Donnelly (2002)). It still gives consistent and asymptotically normal estimates, but the errors calculated from the curvature of the maximum-likelihood surface at its maximum (Fisher-Information matrix) will be too tight when observations are correlated (Lindsay 1988; Coffman *et al*. 2016).

Moreover, in many cases correlations among blocks can be expect to remain fairly weak: Initial correlations of genomic regions in spatial movement are broken up quickly by recombination. When analyzing well separated samples, sharing of long blocks is a very rare event, so most of the observed block-sharing will in fact have independent origin stemming from independent coalescent events. Speaking figuratively, observed IBD-sharing represents the top of whole mountain ranges of underlying correlations, but given that only singular, isolated islands in a sea of improbableness peak out, the massifs of correlation remain unobserved.

#### Poisson model

This assumption of independence is equivalent to modeling the number of shared blocks *k* of a given length between pairs of individuals as independently **Poisson** distributed around the expected rate *λ*:

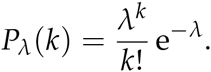

For every pair of chromosomes we bin the shared blocks according to their block length into small bins of width ∆*l*. The limit ∆*l* → 0 would yield exact formulas for block sharing of known lengths, but to have a framework which can deal with errors in block detection it is numerically advantageous to work with finite small bins. In a small bin [*L*_*i*_, *L*_*i*_ + ∆*L*] the expected number of shared blocks can be calculated via Eq. 8:

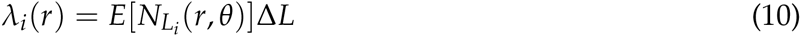

Continuing to assume independence, the likelihood of a pair of samples (*j*) at distance *r* sharing blocks of length 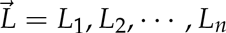 falling into a set of bins *i*_1_, *i*_2_, …, *i*_*n*_ is then given by: 
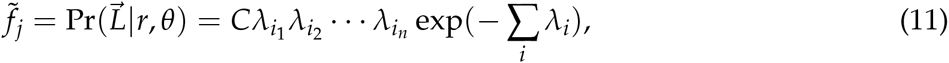
where *C* absorbs all constants that do not depend on the model parameters *θ*- this constant can be dropped when doing likelihood based analysis. Continuing to assume independence, we take the product over all pairwise likelihoods 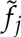 to get the total composite likelihood: 
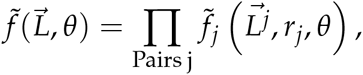
where 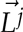 denotes the shared blocks of the jth pair.

The number of pairs 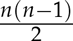 grows quadratically with sample size *n,* and so does the number of terms in the total likelihood. This is advantageous for an inference scheme, but implies that the runtime scales with the square of sample size as well. However, algorithms to maximize functions with a low number of parameters are very efficient, so even sample sizes of hundreds of individuals can be easily handled. Calculation can be also sped up by grouping pairs with the same pairwise distance – such as when analyzing multiple individuals from a population with the same spatial coordinates – since then the *λ*_*i*_ do not have to be calculated repeatedly for every individual pair. Denoting the length bins of blocks shared over all pairs by *i*_1_, *i*_2_ …, *i*_*n*_ and the number of pairwise comparisons by *k* yields: 
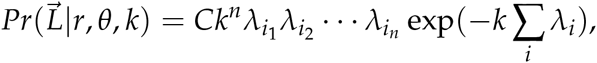
where the factor *Ck*^*n*^ does not depend on the model parameters and can be dropped when maximizing the likelihood.

#### Block detection errors

IBD-block detection is a non-trivial task, and in practice one has to deal with erroneous detection (Browning and Browning 2012). Blocks might be called in the absence of true IBD-blocks (false positives) and on the other hand only a fraction of true IBD-blocks of a given length are detected (limited power), and there is a probability of assigning them the wrong length (error). Following Ralph and Coop (2013) it is straightforward to include these errors into the likelihood framework outlined above. The probability density 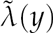 of actually observing a pairwise shared block of length *y* can be calculated from the theoretical probability 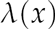 of sharing true blocks of length *x*: 
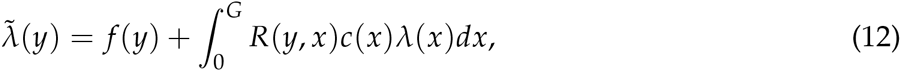
where *f*(*y*) describes the false discovery rate function depending on block length *y, c*(*x*) the power to detect a block of length *x* and *R*(*y*, *x*) the probability of detecting a block of true length *x* as block of length *y*. Doing a careful analysis using techniques such as manually inserting shared blocks and rerunning the IBD-detection allows one to fit these error functions (Ralph and Coop 2013).

In the likelihood framework, for every block length bin of a pair of samples first the predicted true sharing λ is calculated for a set of demographic parameters *θ*, and then updated according to Eq. 12 with the detection error estimates to get the final predicted rates 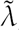, from which the likelihood of observed block sharing can be computed as before. This error model is straightforwardly included into the here presented framework of working with small length bins (Fig. 4).

**Figure 4.**
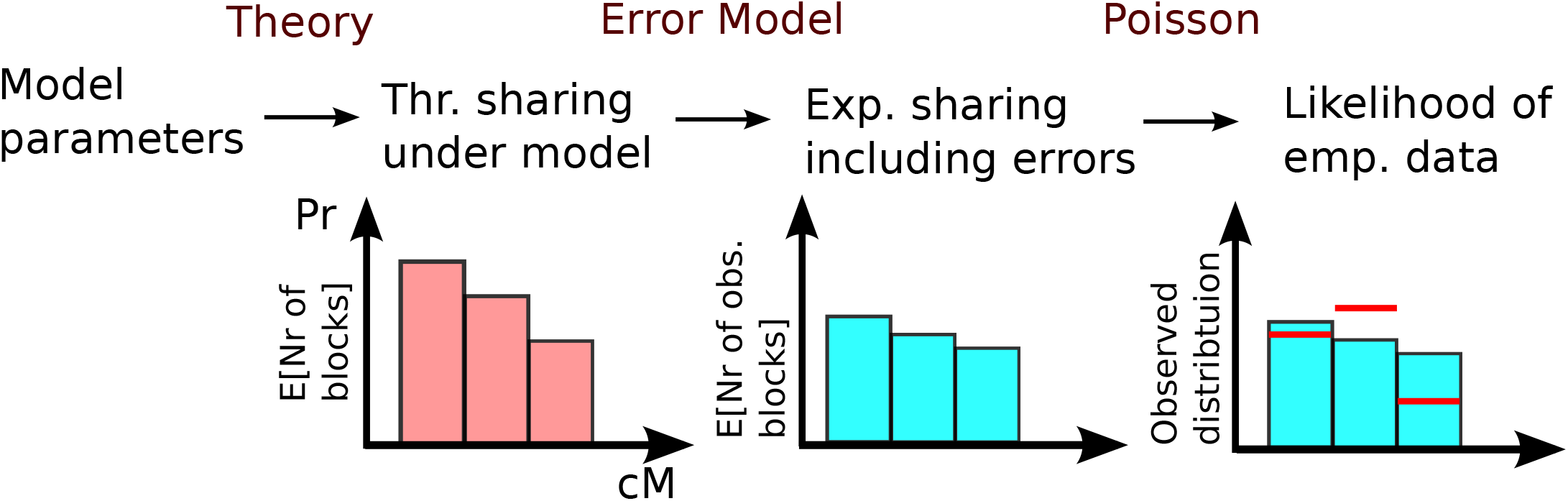
Sketch of dealing with erroneous IBD-data following Ralph and Coop (2013).

#### Adjacent IBD-blocks

The theory and inference scheme introduced here are based on IBD-blocks defined to be ended by any recombination event on the path to the common ancestor. However, multiple consecutive IBD-blocks of recent co-ancestry for unphased data from all four possible pairings of two sets of diploid chromosomes, produce an unbroken segment of exceptionally high similarity which is detected as a single long IBD-block in practice. This can significantly inflate the number of observed IBD-blocks of a given length beyond the true value, especially for shorter IBD-segments (Chiang *et al.* 2016).

The error estimation model of Ralph and Coop (2013) based on introducing artificial IBD-segments of known length can partially account for this, assuming that adjacent IBD-blocks happen to be neighbors by chance, such as when residing on different pairs of homologous chromosomes. However, this does not estimate the effect of short inbreeding loops, where ancestral genetic material broken up by a recombination fuses together again quickly (Fig. 1), rendering the IBD-block ending recombination event "ineffective" (Barton *et al.* 2013).

Intuitively, the larger the neighborhood size (a parameter proportional to the product of *σ*^2^ and effective density: 4*πσ*^2^*D*_*e*_), the smaller the chance that such short inbreeding loops occur. For populations with a large neighborhood size the chance of extending a recent IBD-block by a non-ancient re-coalescence becomes negligible, and our approach remains valid. For instance, simulations of realistic human demography scenarios have demonstrated that the detected number of IBD-blocks is only marginally increased for blocks longer than few cM considered here (Chiang *et al*. 2016).

However, for a population with low neighborhood size, ineffective recombination events can potentially confound observed block sharing patterns and estimates based on it, and unfortunately one cannot hope to account for this within a framework of diffusion of ancestry. This effect is driven mostly by coalescence within few generations, before blocks migrate away from each other. Hence local dispersal and breeding patterns are important; but diffusion of ancestry usually only becomes accurate on intermediate timescales. Therefore, here we empirically study the order of magnitude of this effect by means of simulating specific scenarios with low neighborhood sizes.

### Simulations

To test the above formulas and inference scheme, we simulated sharing of IBD-block in a set of samples by tracing the ancestry of chromosomes backwards in time. Simulations were done on a two-dimensional torus large enough so that IBD-sharing over more than half of the torus was very unlikely, thus effectively simulating a two-dimensional populations without boundary effects. Since sharing of long IBD-blocks is very unlikely to originate too far back in time, we ran the simulations up to a maximal time *t*_*max*_. Errors in block detection are complex and depend on many parameters, and also can be partially accounted for in the estimation framework as described above, so if not otherwise stated, we analyzed sharing of true IBD-blocks, i.e. every recombination event was assumed to be effective.

We used various standard spatial population genetics models:

#### Stepping stone models

In Wright-Fisher stepping stone models every individual occupies a fixed position on a grid and every generation back they choose a parent with probabilities according to a pre-specified backward dispersal kernel. Here, every node is occupied by one or more pairs of two homologous chromosomes, mimicking diploid individuals. For each chromosome a parent position is chosen according to the dispersal kernel and Poisson recombination events along the chromosome induce a switch between the parental chromosomes. Whenever the ancestral material of two distinct initial chromosomes falls on the same chromosome and overlaps for longer than a given threshold chromosome length, we store the resulting IBD-block. We simulated dispersal following discretized uniform, Gaussian and Laplace probability densities along each axis to have representatives of dispersal kernels with low, intermediate and high kurtosis. This model is also easily modified to simulate a nearest neighbor migration deme model: Following pre-specified probabilities a chromosome either chooses its parent uniformly from within its own or one of the neighboring demes.

For simulating the effects of growing or declining population density, we let the number of individuals per node vary with time. A chromosome then first picks an ancestral node as before, and subsequently a random diploid ancestor from this node.

#### Continuous model: Spatial Lambda-Fleming-Viot process

We additionally simulated a model in which each individual occupies a position in continuous space. For this we utilized DISCSIM, a fast implementation (Kelleher *et al*. 2014) of the recently introduced spatial Lambda-Fleming-Viot process. Summarizing briefly (for details see Barton *et al*. (2010)), following lineages backwards in time, events are dropped randomly with a certain rate parameter and uniform spatial density. In each such event, every lineage within radius R is affected with probability u by this event. A pre-specified number of parents, here two, are dropped uniformly within the disc and every affected lineage jumps to them, switching parents according to the recombination rate. Given an initial set of loci, DISCSIM generates their coalescence tree up to a specified time. The output contains a list of all coalescent nodes, which we here further analyzed to detect IBD-sharing.

### Application to Eastern European data

Detecting long IBD-blocks requires population genomic datasets and as of now, such dense genotype or whole genome data of many individuals is available mainly for humans. In order to test the inference scheme, we applied it to a dataset of block sharing between Europeans, which has been generated previously by Ralph and Coop (2013), and includes detailed error estimates for IBD-block detection. They reported significant differences in patterns of block-sharing between eastern and western European populations. Therefore, here we concentrate our analysis on block sharing in the Eastern European subset, as diffusion should be a better approximation for modeling the spread of ancestry in continental regions. Moreover, Eastern European countries are on average geographically more compact, thus position data on country level is expected to be more accurate.

#### The data

The calling and the detailed error analysis of the IBD-data is described fully in Ralph and Coop (2013). Summarizing briefly, IBD-blocks were called for a subsample of the POPRES dataset (Nelson *et al*. 2008) genotyped at ~500, 000 SNPs using the fastIBD method, as implemented in Beagle v3.3 (Browning and Browning 2011). Every sample used in the analysis was required to have reported grandparents all from a single country. Here we analyze block sharing between 125 Eastern European samples (see SI Text 1). We follow the geographic classification of (Ralph and Coop 2013), but exclude the six Russian and one Ukrainian samples, as location data at country level is likely very inaccurate for these two geographically extended countries We analyzed shared blocks longer than 4 cM, and within our subsample 1824 pairwise shared blocks have been reported. We set the position of each country to its current demographic center, defined as the weighted mean location (SI Text 1); and calculated pairwise distances based on this.

#### Data analysis

Throughout the analysis, we worked with block length bins ranging from 0 to 30 cM with a bin width of Δ*l* = 0.1 cM and applied the error function estimates reported by Ralph and Coop (2013) as described above. For maximizing the likelihood, we calculated the likelihood of block sharing in the bins from 4 to 20 cM, which is informative about the last few centuries (see Fig. 11). We excluded longer shared blocks from our analysis, since these have a considerable chance of originating in the last few generations, which is not expected to be accurately captured in the diffusion model and is moreover also confounded by the sampling scheme that always requires four grand parents from the same country.

Within the likelihood framework described above, we fitted several specific models of past density *D*:

- A constant population: *D* = *C:*
- A population growing at accelerating rate: *D* = *C/t*
- A growth model where the growth rate is fitted as well: D = *Ct^−β^,*

where *t* measures time back in generations. To learn about the certainty of estimates, in addition to using the curvature of the likelihood surface (Fisher-Information matrix) we bootstrapped the data. Since we suspect strong correlations and systematic deviations from the model, we re-sampled over different units: We bootstrapped on block level by redrawing each block a number of times following a Poisson distribution of mean 1, and similarly over country pairs, since we suspect systematic correlations for certain population pairs.

Furthermore we analyzed the residuals from the best fit models, i.e. the deviation of pairwise block sharing between pairs of countries from the expected value predicted by the best fit model. For this we assumed that the observed block sharing to be Poisson distributed around the predicted block sharing. Transforming the block count data 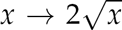 transforms Poisson distribution into approximately normal distributions with standard deviation 1, which aids visual inspection of the statistical significance of residuals.

### Data Availability

We implemented the above described methods to simulate and analyze IBD-sharing data in Python.

The source code has been uploaded to the freely available Github repository https://git.ist.ac.at/harald.ringbauer/IBD-Analysis.

The preprocessed human IBD-block sharing data including detection error estimates used here are the result of the analysis of Ralph and Coop (2013) and can be freely accessed at http://www.github.com/petrelharp/euroibd.

## Results

### Block-sharing in simulated data

Here we present results for block sharing from simulated data and compare them to theory. When visualizing the results we always bin IBD-block sharing with respect to block length and sample distance and depict rates per pair for the bin, and normalize for a rate per cM.

#### Constant population density

For the important case of constant population density we compare simulated IBD-sharing to the theoretical expectation (Fig. 3) as calculated from Eq. 8:

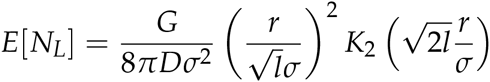

Here *K*_2_ denotes the modified Bessel-function of the second kind of degree 2 (Abramowitz and Stegun 1964). This formula predicts that block-sharing approaches exponential decay with distance, as Bessel-functions 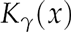 converge to 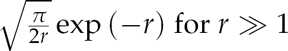 (Abramowitz and Stegun 1964). This decay then dominates the polynomial terms in front of the Bessel-functions and the slope of this exponential decay (on a log scale) approaches 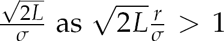. For the long blocks considered here, this happens already within few dispersal distances. In all simulations, block-sharing patterns were very similar among the five different simulated models, and closely followed the theoretical expectation.

#### Growing and declining populations

In Fig. 5 the results of simulations for three population density scenarios are compared to the predictions of Eq. 8. Here we depict the result for simulated Laplace dispersal, the other dispersal kernels yield similar results. Again, simulation results are in close agreement with theory. The decay of block sharing with distance approaches exponential decay with rate 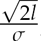, where the specific density scenario determines the speed of convergence - Bessel functions *K*_*γ*_ approach the exponential regime slower with growing *γ*.

**Figure 5.**
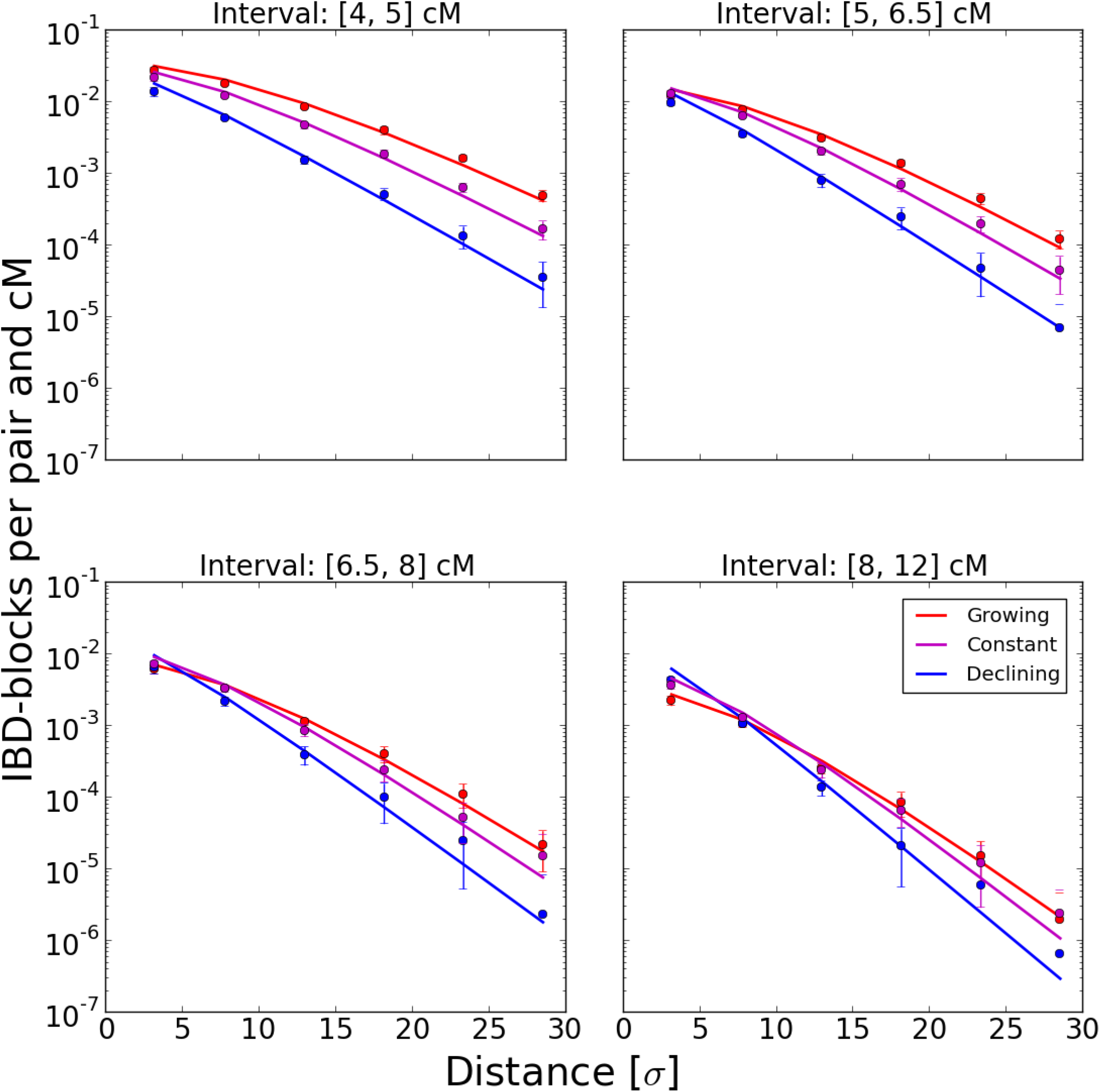
Various population density scenarios: Simulated IBD-block sharing per pair and cM in various density scenarios compared to Eq. 8: Block sharing of a subset of 150 cM chromosomes 4 distance units apart placed on an initial grid was analyzed. Along each axis dispersal was modeled by a Laplace distribution with *σ* = 1 and the number of diploid individuals per node *n* either remained constant at *n* = 10, grew as *n*(*t*)= *t* or declined as *n*(*t*)= 200/t where in all cases *t* denotes the time back measured in generations and at every step *n*(*t*) was rounded to the nearest integer value. For each scenario 20 replicate runs were done, dots depict the mean and error bars the standard deviation for every bin, line the predicted values from Eq. 8.

### Inference in simulated data

We tested the parametric inference scheme introduced above and analyzed its ability to recover the underlying demographic parameters from simulated block sharing data. For every simulated block data set, we numerically computed the maximum likelihood estimates *θ*_*MLE*_ as described above for shared blocks between 4 and 20 cM, put into bins of width 0.1 cM. After testing multiple methods we decided to use the Nelder-Mead method here, as implemented in the class GenericLikelihoodModel of the Python package statsmodels, because it proved to be numerically stable and quick. We also estimated standard deviations and confidence intervals from the empirical Fisher-Information matrix by using implementations from the same package.

#### Constant Population Density

Results for parameter inference for a population of constant density are depicted in Fig. 6. We simulated a varying number of samples on a grid, consisting of one chromosome each, to test the behavior of the inference scheme with respect to limited sample size. Naturally, the variance of estimates increases with decreasing sample size, but the bias remains small. Moreover the estimated standard errors are capturing the true estimator variance relatively well (see SI Text 2). This confirms that most of the shared blocks are the result of uncorrelated coalescence events, as heuristically argued above. The typical log-likelihood surface for a single simulated IBD-block sharing data set is found to be smooth (Fig. S2), and in all cases numerical maximization did not result in spurious maxima, even for initial estimates orders of magnitudes off. Moreover, estimates of density and dispersal rate are only slightly correlated in the scenario considered here (SI Text 2).

**Figure 6.**
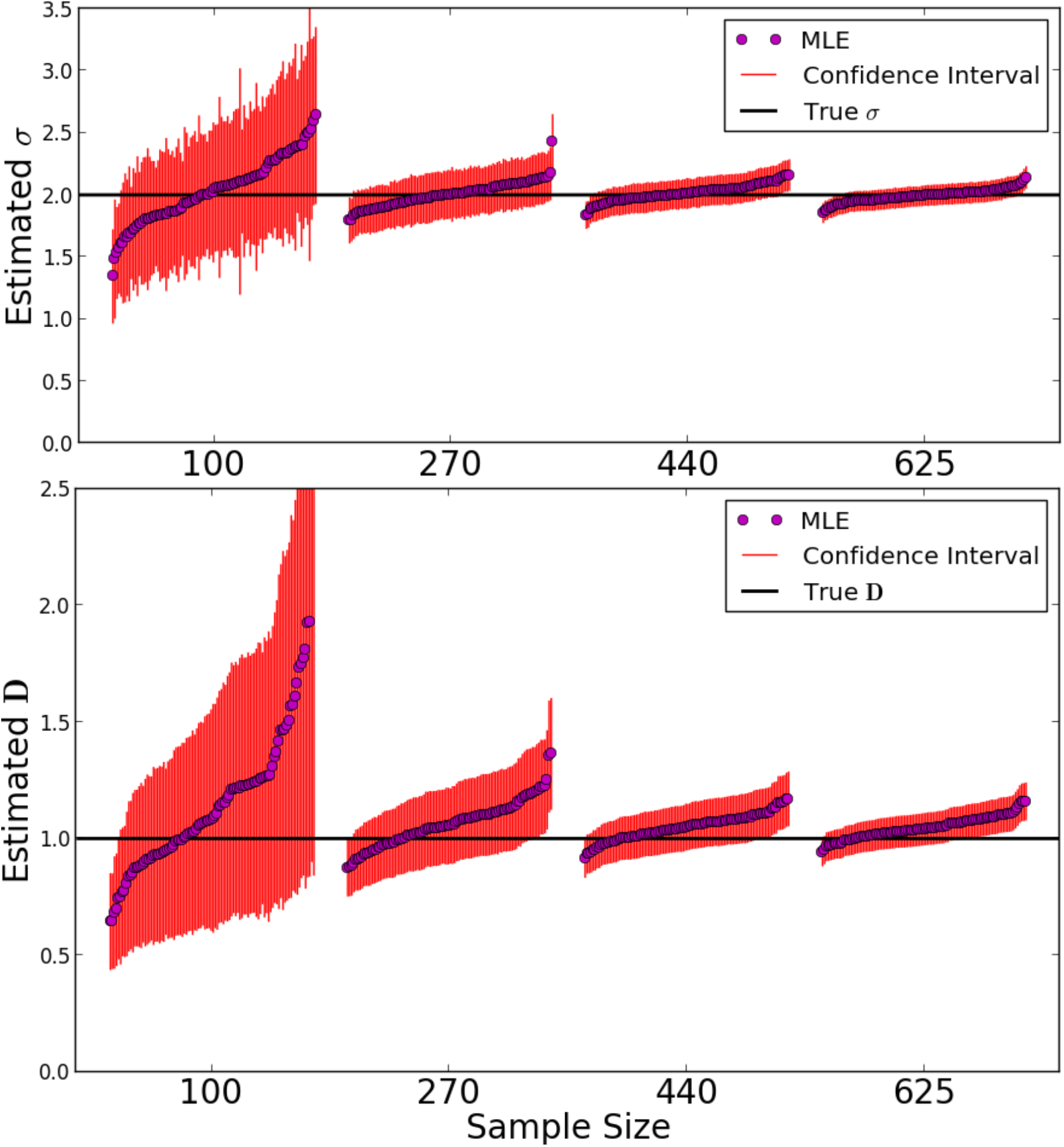
Maximum Likelihood estimates: We simulated a Laplace model on a grid of nodes of torus length 180 for *t* = 200 generations back. We set the dispersal rate *σ* = 2 and the number of individuals per node to *D* = 1. In every run, a random subset of 150 cM chromosomes was picked from a initial sample grid spaced two nodes a apart to reach one of four sample numbers: 100, 270, 440, 625. For each sample size 100 simulations and subsequent parameter estimates from the IBD-data were run. Every dot depicts the MLE parameter estimate of a single run and the 95% confidence intervals calculated from the Fisher-Information matrix.

#### Varying Population Density

We also tested the ability of the inference scheme to detect recent changes in population densities. For this we simulated three scenarios of a growing, declining and constant population with growth parameters *β* = 1,0, –1. Results are depicted in Fig. 7. The estimates of the demographic parameters allow one to robustly distinguish these three scenarios. Interestingly, accurate estimates of the dispersal rate are feasible in all these demographic scenarios; even when fitting a model with constant population size to the other two scenarios of a recently quickly changing population size (Fig. S1). This can be heuristically explained by the fact that the eventual rate of decay, the main signal for estimating *σ* from fitting Eq. 8, remains the same, independent of the specific population density scenario. The speed of convergence varies, but in all cases the eventual rate is approached relatively quickly within several dispersal distances (Fig. 5).

**Figure 7.**
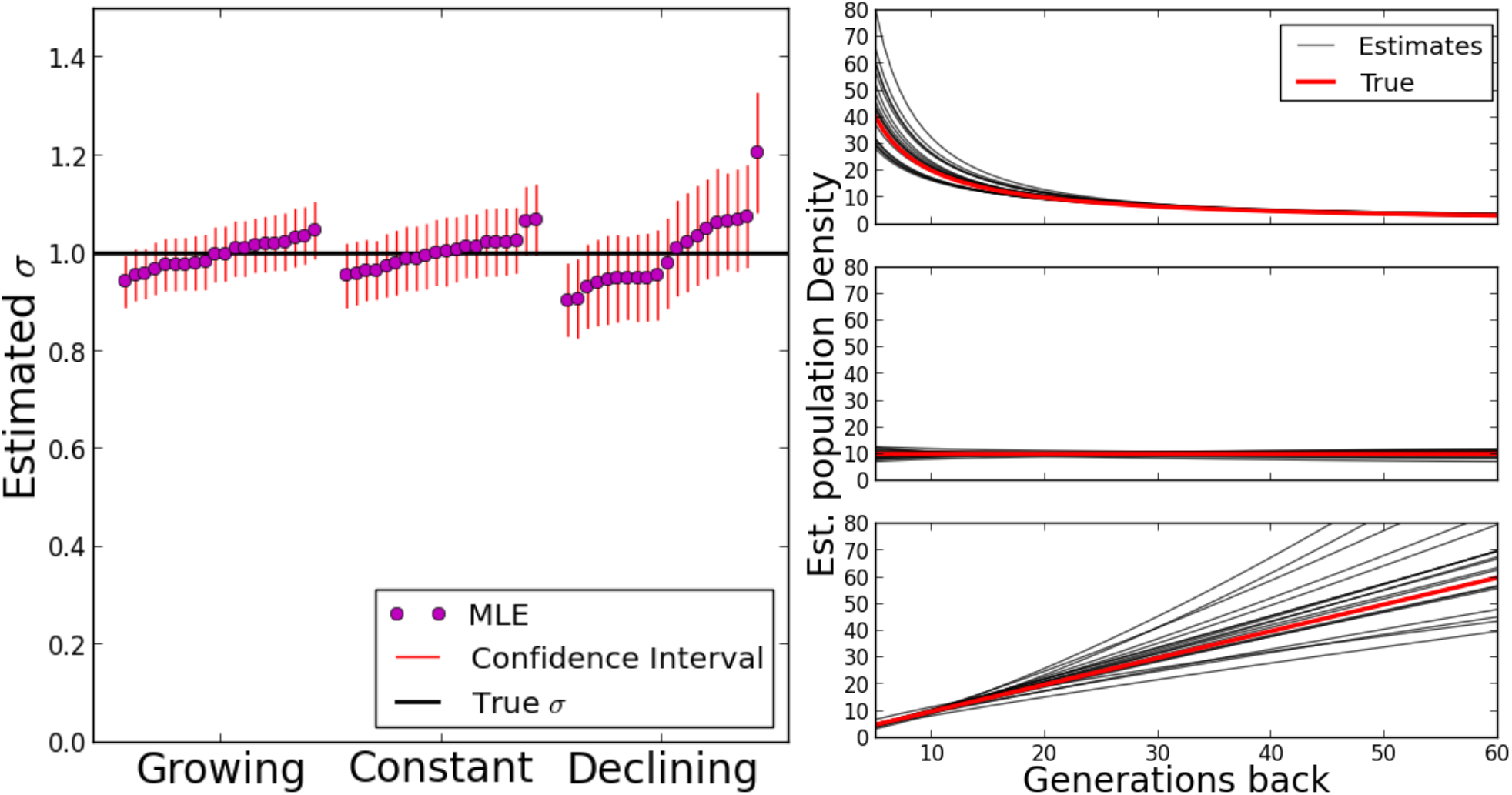
Likelihood estimates for various population density scenarios. The same scenarios as in Fig. 5 were simulated. For 20 runs each, 625 chromosomes of length 150 cM were randomly picked from a sample grid and traced back using Laplace dispersal with *σ* = 1, and maximum likelihood fits and 95% confidence intervals calculated from their block sharing. For the estimated population density the true value of simulations and the MLE-estimate for every run are shown.

#### True versus detectable IBD-blocks

In Fig. 8 the above described effect of undetected recombination events on estimates of demographic parameters and overall IBD-block number is depicted. This was investigated with simulations in the DISCSIM model, as it allows easy and continuous tuning of the neighborhood size *4πσ*^2^*D*_*e*_ via the parameter describing the probability that an event hits an individual within its range (Barton *et al.* 2013). It is also computationally very effective: the complete ancestral material of all initial samples can be traced back without the need to delete short blocks. Pairwise coalescence times for all pairs of loci along the chromosome were extracted, but now, only effective recombination events are counted, defined here as jumps of coalescence times between adjacent pairs of loci with at least one coalescence time older than a preset time threshold of 1000 generations back, the time until which the backward simulations were run here. This should capture most short recombination-coalescence loops while still being well below the bulk of ancestral coalescence times. The effect of non-detectable recombination events becomes significant only for very low neighborhood sizes (< 15), when the detected number of IBD-blocks of a certain length gets inflated by wrongly inferring multiple shorter blocks as a single longer block. While estimates for density remain almost unbiased, the inferred dispersal rates increase significantly, likely explained by an excess of block sharing for distant samples. However, even for very low neighborhood sizes, when the density of individuals measured in dispersal units (*σ* = 1) is about one (for neighborhood size ≈ 4), the upward bias remains less than 50%.

**Figure 8.**
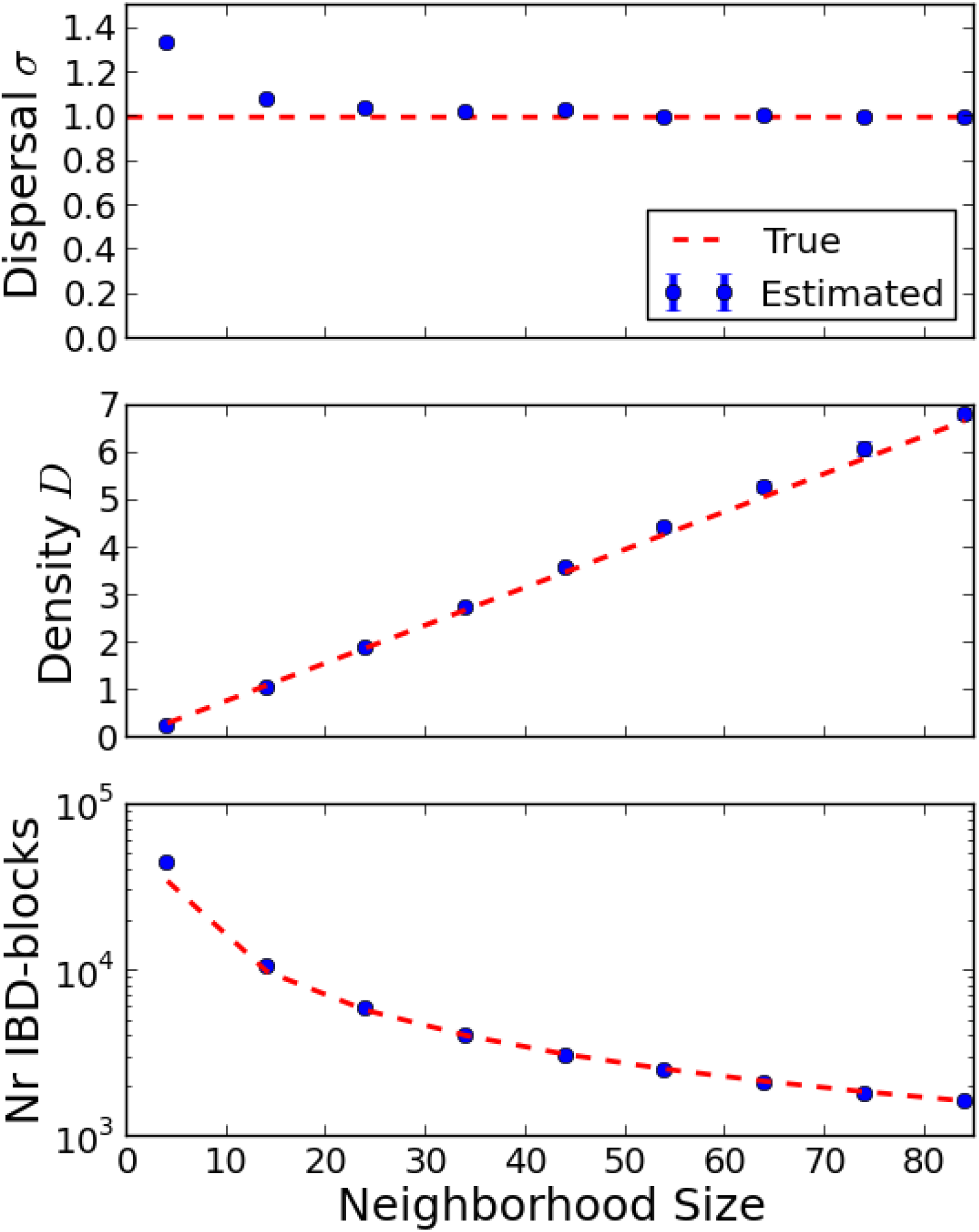
Observable IBD-blocks and estimates vrs theoretical prediction for true IBD-blocks. Simulation were run with DISCSIM for an initial grid of 150 cM chromosomes three distance units apart on a torus with axial size 90. Dispersal rate was set to 1. IBD-blocks were detected as consecutive runs of coalescence times < 1000 generations and then parameters inferred as described above. For various densities, corresponding to neighborhood sizes 4 - 86, ten DISC-SIM runs were simulated. The mean of these runs is compared to calculations based on theory for true block sharing (Eq. 8).

### POPRES data

#### Best fit models

When fitting our models to the Eastern European subset of the POPRES IBD-data, the model of quick population growth with a population density 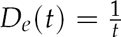 for *t* generations back fits markedly better than a model of constant population size, which underestimates sharing of short blocks (Fig. 9) at its maximum likelihood parameters. The more complex model *D*_*e*_(*t*) = *t*^*-β*^ where the growth rate parameter *β* is estimated as well estimates *β* to be close to 1 and the increase of likelihood is small (Δ*L* = 1.1), especially when considering that there are correlations in the data which make the difference of true likelihood even smaller (Coffman *et al.* 2016). Similarly, fitting several more complex density functions as sums of power terms did not significantly increase the likelihood. In all three models, estimates for dispersal *σ* are about 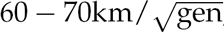, even under the likely misspecified constant population size model (Tab. 1), and bootstrapping on country pair level yields 95% confidence intervals ranging from 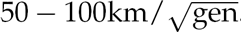.

**Figure 9.**
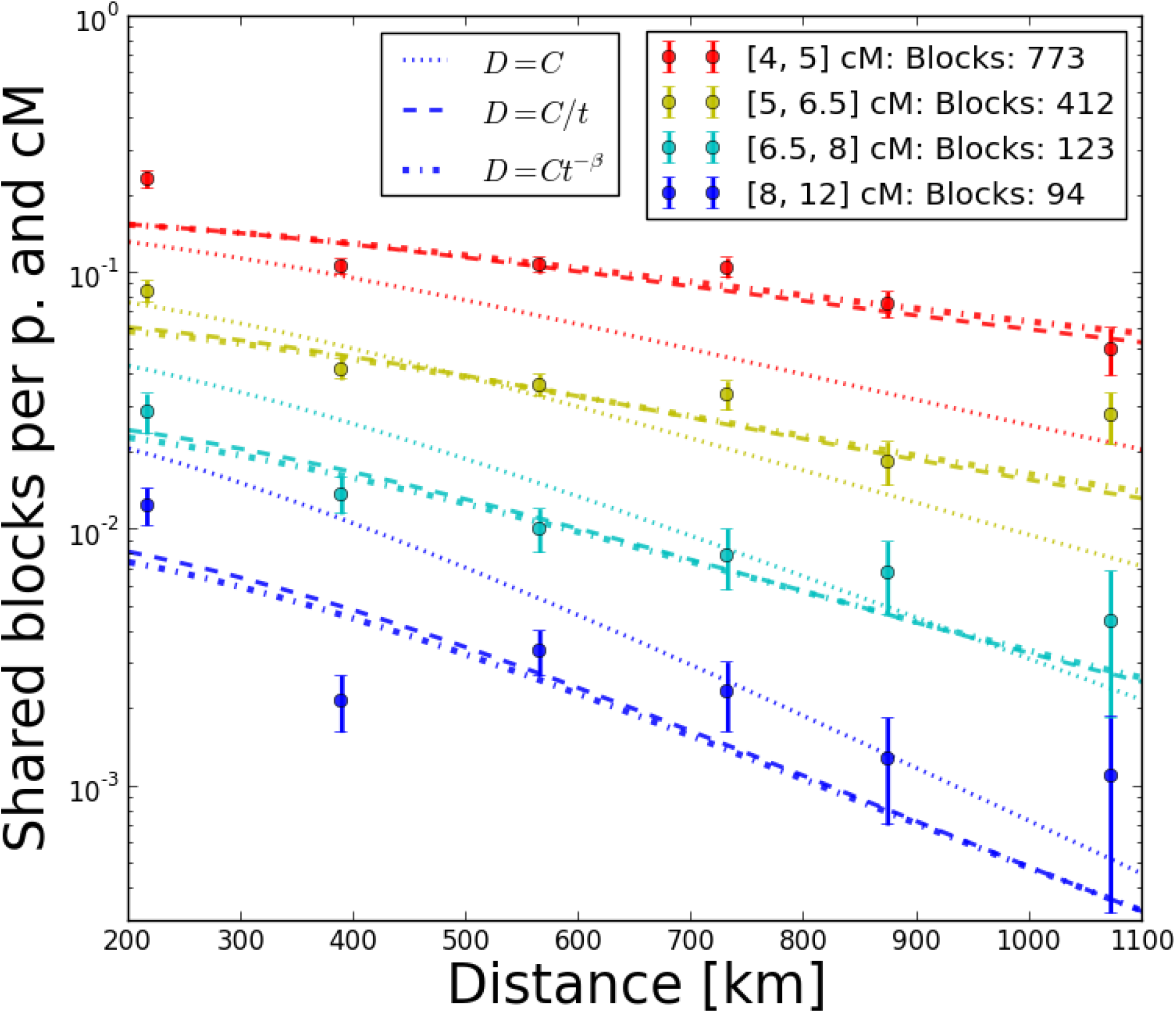
Fit of three models to Eastern European block sharing data. Dots depict mean block sharing within distance and block length bins for blocks of several length categories, the lines are prediction for the best fit models. The error-bars represent standard deviations under the assumption of Poisson counts in every bin – some are clearly too tight and there are outliers, which hints at more systematic deviations at country-pair level (see also Fig. 10)

**Table 1.** Maximum Likelihood estimates for Eastern European IBD-data.

| Density Model | Parameter | MLE-Estimate<sup>a</sup> | 95% CI<sup>b</sup> | 95% CI Bootstr.<sup>c</sup> |
| --- | --- | --- | --- | --- |
| $D_e = D$ | $D$ | 0.047 | 0.043 - 0.051 | 0.038 - 0.065 |
| | $\sigma$ | 67.8 | 62.9-72.8 | 53.03 - 81.50 |
| $D_e = D/t$ | $D$ | 1.71 | 1.48-1.94 | 1.22 - 2.87 |
| | $\sigma$ | 62.6 | 56.1-69.0 | 42.2 - 82.6 |
| $D_e = Dt^{-\beta}$ | $D$ | 2.13 | 1.39-2.86 | 1.16-5.83 |
| | $\sigma$ | 63.0 | 56.2-69.8 | 44.2-82.4 |
| | $\beta$ | 1.05 | 0.98 - 1.13 | 0.90-1.25 |

<sup>a</sup> All units so that distances are measured in km and time in generations
<sup>b</sup> Calculated from Fisher-Information Matrix
<sup>c</sup> Calculated from 100 estimates on data bootstrapped over country pairs

The estimated parameter uncertainty when bootstrapping over single blocks is only slightly bigger than estimated from the curvature of the likelihood, but bootstrapping over country pairs gives markedly increased confidence intervals (Fig. S3). This implies that there are strong correlations at country pair level in the data. This is further confirmed by the analysis of residuals for country pairs, which yields a gradient towards the Balkans for more block sharing than predicted by the best fit models. This is statistically most significant for short blocks due to increased power due to the higher number of shared blocks (Fig. 10), but the overall pattern holds also for longer blocks (Fig. S4).

## Discussion

### Exponential decay of IBD-sharing with distance and block length

The derived formulas for sharing of long IBD-blocks under diffusion of ancestry are structurally similar to the Wright-Malecot formula (Barton *et al.* 2013) that describes allelic identity by state using similar approximations: A polynomial factor is multiplied with a Bessel-Function of the second kind, 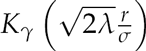, where λ is the rate of the identity destroying process. For allelic correlation λ = *µ*, the mutation rate, while here *λ* = *L*, the IBD-block map length. For long blocks L is much bigger than typical mutation rates *µ*. This allows one to probe the tail of the Bessel-function 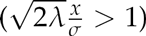 where it approaches exponential decay that dominates the polynomial factor. This exponential decay occurs with both increasing block length 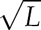 and importantly increasing geographic distance *r,* where usual a wide range of different sampling distances are available. Theory predicts that for long blocks this decay can be over orders of magnitudes for multiple dispersal distances, which is observed in the human data (see Fig. 9, 3). This pattern persists even in the case of relatively quick recent population density change (see Fig. 5). When global density changes can be modeled as sums of power terms of the form Eq. 7, the result for expected block sharing will be given by sums of corresponding Bessel-Functions (Eq. 8). Each of those approaches exponential decay, and so also their sum does. Thus, when density change is slower than the quick process breaking up blocks, theory predicts that the exponential decay with distance will generally approach an eventual rate 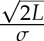. Therefore, estimates of the dispersal rate *σ* making use of the decay rate in the exponential regime can be expected to be relatively stable with respect to recent demographic history (see also Fig. S1).

**Figure 10.**
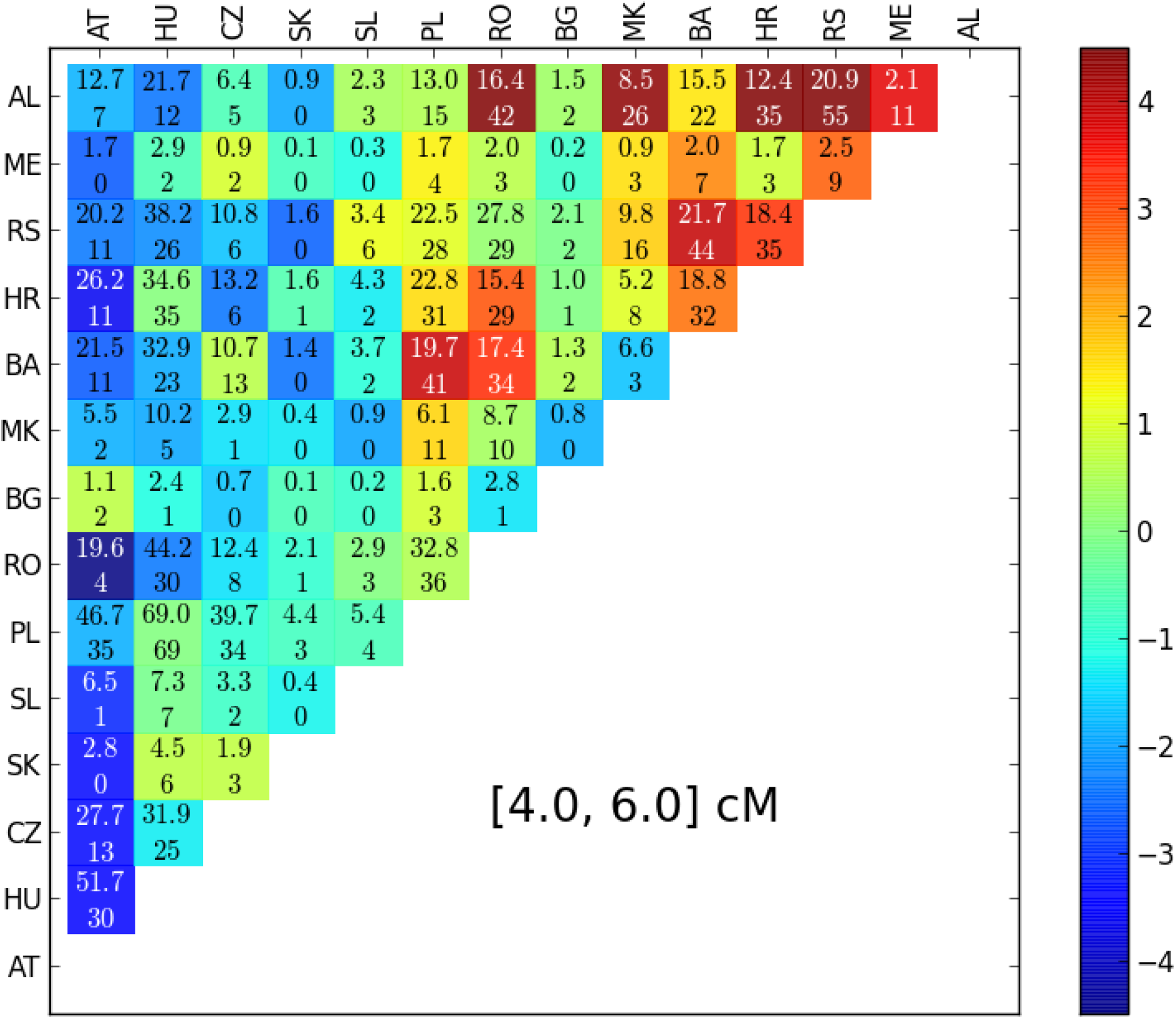
Residuals for pairs of countries for blocks of length 4 – 6 cM: Upper line in every field: Total number of IBD-blocks expected from best fit model and sample size. Lower Line: Observed number of IBD-blocks. Color of every field is determined by statistical significance (z-Value when transformed 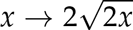. (AT: Austria, HU: Hungary CZ: Czech Republic, SK: Slovakia, SL: Slovenia, PL: Poland, RO: Romania, BG: Bulgaria, MK: Macedonia, BA: Bosnia, HR: Croatia, RS: Serbia, ME: Montenegro, AL: Albania)

**Figure 11.**
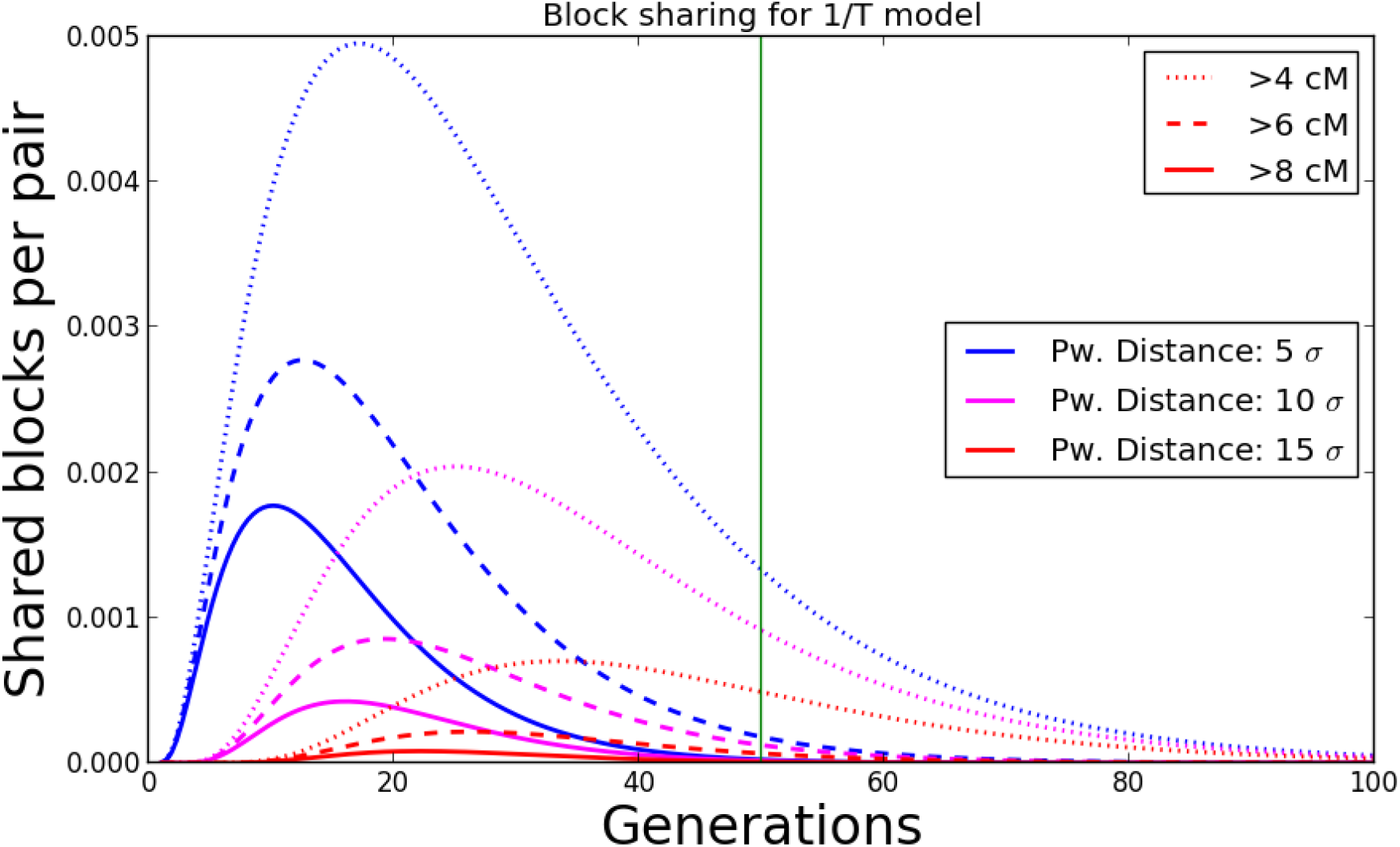
When do the observed IBD-blocks come from? Density of blocks of certain length originating *t* generations ago, as calculated from the 1/*T* population density growth model with best fit parameter for density (best fit 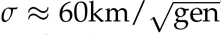). Most of the signal is predicted to have arisen within the last 50 generations (green line)

### Implications for demographic inference

The fast rate at which long blocks are broken up and the ability to probe the exponential regime of decay offers several significant advantages for demographic inference. First, long blocks typically stem from very recent time (see Fig. 11). This is clearly advantageous for populations which have been in equilibrium for only relatively short time – as is likely often the case. Inference methods that rely on allelic correlations do probe very recent time scales as well (Barton *et al*. 2013), in fact they pick up increased identity by state induced by the here described shared blocks, but this is on top of a majority of identity by state stemming from more ancient times. Thus these methods are much more susceptible to confounding by ancestral structure (Meirmans 2012) and have stringent, often unrealistic, equilibrium time requirements (Leblois *et al*. 2003), which the here introduced method can avoid. For instance, in the human case the best fit model predicts that most long blocks stem from within the last 50 generations (Fig 11). Second, quick exponential decay, both with sampling distance and also block length, offers a very robust signal for demographic inference. As demonstrated here, sharing of a few cM long blocks decays quickly over orders of magnitudes within several dispersal distances and estimating rates of such quick decay is very stable, as demonstrated by the accuracy of the inference method on simulated data. This is in contrast to inference based on allelic correlations, which only decay with the logarithm of distance (Barton *et al.* 2002). Indeed, studies using allelic correlations report low and problematic signal to noise ratios (Watts *et al.* 2007). Third, as mentioned above, utilizing the logarithmic regime of the Wright-Malecot formula only allows one to infer the neighborhood size, the product of density and dispersal 4*πDσ*^2^. Naturally, however, often their separate values are of interest. As demonstrated here, using long blocks and recombination as a quick clock with known rate allows one to obtain separate estimates of the two.

The results here show that sharing of long blocks can decay over orders of magnitudes within as little as a few dispersal distances. This urges caution when fitting block sharing to simple models of panmictic populations to learn about recent population sizes (for instance (Browning and Browning 2015)), as local population structure can heavily influence IBD-sharing patterns. If assumptions about panmixis do not hold and sample location influences IBD-sharing patterns, results should not be interpreted in terms of recent population size. In contrast, the method introduced here can estimate effective population density and global patterns of population growth or decline by explicitly taking spatial sample locations and spread of ancestry into account.

The here used inference scheme is based on a simple model of diffusion of ancestry. While reality will be often more complex, having a simple model that yields analytical formulas is a big computational advantage for doing demographic inference. The predicted quick decay of block sharing both with distance and also block length offer a stable signal for inference, and so even in face of complex model deviations, estimates for ecologically and evolutionary important parameters such as the typical dispersal rate should be relatively robust.

### Analysis of human data

The analysis of human data nicely demonstrates our inference scheme. The true demographic scenario is doubtless more complex, including heterogeneous, time-dependent migration rates and large scale migrations. That said, the broad patterns of IBD sharing can be fitted well with a diffusion model. Using our inferred model we predicted most of the shared blocks we use (> 4 cM) and hence our signal originates within the last 50 generations (Fig. 11), corresponding to the past 1450 years (assuming 29 years per generation (Fenner 2005)). This mostly postdates the period of large scale migrations in Europe ("Völkerwanderung" (Davies 2014)). Our inferred demographic parameters seem to be plausible: There is a clear signal for rapidly accelerating recent population growth, which is in a agreement with historical estimates (McEvedy *et al.* 1978) and previous genetic studies based on the allele frequency spectrum (Keinan and Clark 2012; Gao and Keinan 2016). The inferred dispersal rate 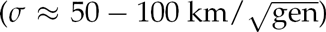 also agrees with historical studies, which imply that preindustrial individual human migrations over dozens of kilometers are rare but happen at a significant rate (Wijsman and Cavalli-Sforza 1984; Pooley and Turnbull 2005).

Here we also detect a systematic, large scale deviation from a simple diffusion model with uniform population density, as there is a clear gradient for higher block sharing towards Balkan countries (Fig. 10), which was already observed by Ralph and Coop (2013). They hypothesized that this could be due to the historic Slavic expansion, a hypothesis supported by admixture analysis (Hellenthal *et al*. 2014). However, the pattern of increased block sharing also holds for longer, typically younger blocks, which could hint additionally at a consistently lower population density in these regions. Such systematic regional deviations from the diffusion model also imply that care should be taken when interpreting parameter estimates and their uncertainty estimates.

### Outlook

The inference scheme based on long IBD-blocks introduced here is data-hungry and requires population genomic datasets, as it relies on dense genotype data from at least a few dozen individuals. It requires spatial information of the samples, and also at least a coarse linkage map. However, the novel opportunities and advantages for inference of recent demography should well justify the effort. The possibility to accurately estimate dispersal distances and past effective population density has the potential to yield interesting novel insights for a whole range of organisms. The necessary datasets are already within reach for several systems, and they will become even more accessible in the near future with increasing genotyping capacities. In cases when there are severe deviations from the model of uniform spatial diffusion of genetic ancestry, the here introduced inference scheme based on analytical formulas will be a helpful first step for identifying those. In order to fully utilize the potential of shared IBD-blocks, future work is needed to develop methods that go beyond inference of the most basic parameters of demography and can fit more complex migration patterns and population structures. We hope that the here introduced inference scheme and the underlying concepts mark only a step in a new era of demographic inference.

## Appendix

## Appendix A

We use diffusion to model the pairwise distance of two lineages backward in time: Let *r*(**x**, *t*) denote the density function describing the probability to have pairwise distance **x** at time *t* back. Coalescence is included into this model by modeling that two lineage coalesce instantaneously at an average coalescence rate *v*(**x**) depending on the pairwise sample spacing **x**. For the probability of coalescing at time *t* ago, we get:

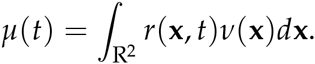

In models where only discrete sample distances **x** are possible, such as the stepping stone model, the integral is replaced with a sum. The key observation is that v(**x**) is usually negligible outside a small area around the origin, since in most models only very close samples (|**x**| ≈ *σ*) have an appreciable chance to coalesce. Within such small areas around the origin, for *t* >> 1 we approximate *r*(***x***,*t*) with ≈ *r*(0, *t*) and get: 
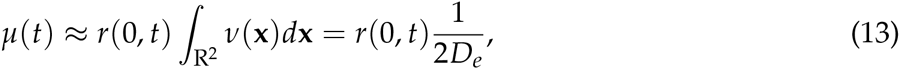
where we have defined 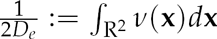. This is the final approximation used for deriving our formulas for block-sharing. It can be shown that stepping stone models asymptotically converge to this when rescaling appropriately (Barton *et al.* 2002, 2013). With demes separated by one distance unit *D*_*e*_ corresponds to the number of diploid individuals per deme, which motivates the name effective density. Here we give this more general definition of *D*_*e*_ to allow one to directly calculate its value in various scenarios we simulated above (see SI Text 2).

## Appendix B

Here we give the full result for block sharing that includes chromosomal edge effects, which we use for inference. We shall denote the formula Eq. 8 with fixed *G* = 1M with *n_L_*(*β*), where the dependencies other than *β* are suppressed for ease of notation. Then, integrating Eq. 3 yields: 
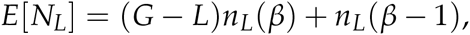
the formula for one chromosome of length *G*. For multiple chromosomes of different lengths one has to sum this formula over all chromosomes. For pairs of diploid individuals, the resulting formula has to be also multiplied by a factor of four, since for every pair of individuals four pairs of chromosomes are compared.

For the analysis of human data, here we use sex average map lengths of autosomes given by the Decode map (Kong *et al*. 2002), consistent with (Ralph and Coop 2013).

